# Channel-mediated astrocytic volume transient is required for synaptic plasticity and spatial memory

**DOI:** 10.1101/2025.02.03.636214

**Authors:** Junsung Woo, Victor James Drew, Jung Moo Lee, Wuhyun Koh, Joungha Won, C. Justin Lee

**Affiliations:** Department of Neuroscience, Division of Bio-Medical Science & Technology, KIST School, Korea University of Science and Technology (UST), Seoul 02792, Republic of Korea; Center for Cognition and Sociality, Institute for Basic Science (IBS), Daejeon, 34126, Republic of Korea

**Keywords:** Astrocytic volume transient, TREK-1, TRPA1, BDNF, synaptic plasticity, spatial memory

## Abstract

Astrocytes, known for their support roles, are emerging as active participants in synaptic plasticity and cognitive functions. Astrocytes actively regulate synaptic plasticity and memory through dynamic volume transients. Our previous research identified several key molecules, including TREK-1, TRPA1, and Best1 ion channels, as well as the gliotransmitter BDNF, as critical components of astrocytic volume transients. However, the precise mechanisms by which these volume transients influence synaptic plasticity and memory remain poorly understood. In this study, we investigate the roles of TREK-1 and TRPA1 in astrocytic volume dynamics and their downstream effects. Using intrinsic optical signal imaging, electrophysiology, and behavioral assays, we demonstrate that neuronal stimulation induces astrocytic swelling, initiated by K^+^ uptake through TREK-1 channels and regulated by Ca^2+^ influx via TRPA1 channels. This swelling is closely associated short- and long-term potentiation, and is accompanied by the release of BDNF, which restores long-term potentiation under conditions of calcium sequestration during astrocytic calcium clamping experiments. Disruption of astrocytic volume transient associated ion channels results in significant deficits in spatial memory, as evidenced by impairments in object-place recognition and passive avoidance tasks. Furthermore, these channels were found to modulate the synaptic plasticity. These findings reveal astrocytic volume transients and BDNF as pivotal modulators of synaptic plasticity and memory, as well as potential therapeutic targets for addressing memory dysfunctions.

## 1. Introduction

Synaptic plasticity, the cellular mechanism underlying learning and memory, is a highly dynamic process orchestrated through intricate interactions between neurons and glial cells.^1–3^ Among these, astrocytes—once thought to only play a supportive role—are increasingly recognized as active regulators of synaptic and circuit function. Astrocytic responses to neuronal activity involve transient volume changes driven by ion influx, accompanied by water entry mediated through water channels such as aquaporin-4 (AQP4).^4^ These volume transients have been implicated in modulating the synaptic microenvironment, potentially influencing plasticity and cognitive function.^5^

Astrocytic volume transients are closely coupled to neuronal activity.^4^ Potassium uptake, facilitated by TWIK-related K⁺ channel 1 (TREK-1)-containing two pore potassium (K2P) channels, triggers water influx through AQP4. This creates a swelling phenomenon that is followed by chloride efflux via bestrophin-1 (Best1) anion channels, enabling volume recovery.^4^ These processes have been linked to synaptic potentiation and memory formation, as demonstrated by recent studies showing impaired long-term potentiation (LTP) and memory in AQP4-deficient models.^6^ Furthermore, the dynamics of astrocytic swelling, captured through intrinsic optical signal (IOS) imaging^4^, provide insight into their contribution to activity-dependent extracellular ion and neurotransmitter homeostasis.^7^ Despite advances in understanding astrocytic contributions to plasticity, several critical gaps remain. While prior research has described the general roles of channels such as TREK-1 and transient receptor potential ankyrin 1 (TRPA1), the specific interplay between astrocytic calcium signaling and the release of gliotransmitters like brain-derived growth factor (BDNF) has not been fully delineated.

Among the gliotransmitters released, BDNF stands out due to its high expression levels in the brain and its well-established potent effects on synaptic plasticity.^8,9^ Pioneering research in the early 1990s demonstrated that a stimulation protocol capable of inducing LTP in the hippocampal CA1 region also increases BDNF mRNA expression in neurons.^10^ Further investigations revealed that LTP is compromised in BDNF gene-silencing mouse models but can be restored by reintroducing BDNF.^11,12^ BDNF is associated with accelerated synaptogenesis in various subtypes of neurons, as well as improved memory function through NMDA receptor signal activation.^13,14^ Interestingly, it has been demonstrated that astrocytic BDNF is also capable of modulating LTP and memory formation.^15^ Notably, the direct influence of astrocytic volume transients and corresponding calcium dynamics on both short-term and long-term plasticity, as well as their implications for spatial memory, remains an area of active investigation.

This study addresses these gaps by examining the hypothesis that astrocytic calcium signaling, mediated through volume transient activation of mechanosensitive channels and BDNF release, is essential for regulating synaptic plasticity and memory. Using a combination of electrophysiology, genetic manipulations, and behavioral assays, we investigate how astrocytic mechanisms impact synaptic potentiation and memory. By employing advanced tools such as tamoxifen-inducible Cre-lox systems^16^ and pharmacological interventions, we provide new insights into how astrocytes support plasticity and memory through calcium-dependent mechanisms and the targeted release of BDNF. Our findings not only highlight the critical functions of astrocytic volume transients and calcium signaling but also establish a mechanistic link between astrocyte-neuron interactions and memory formation, offering a deeper understanding of glial contributions to brain plasticity and memory.

## 2. Methods

### 2.1 Animals

Adult male mice (6–10 weeks old) were used in this study. Wildtype mice (C57BL/6; Jackson Laboratory, RRID: IMSR_JAX:000664), TRPA1 knockout (TRPA1 KO) mice, (129 strain; Jackson Laboratory, RRID: IMSR_JAX:006401), GFAP-GFP (JAX stock #003257) mice and their respective wildtype littermates were included. Mice were housed in groups of 3–5 per cage under a standard 12-hour light/dark cycle with ad libitum access to food and water. All experimental procedures were approved by the Institutional Animal Care and Use Committee (IACUC) of the Institute for Basic Science (IBS, Daejeon, Korea; Protocol No. IBS-22-26) and conducted in accordance with institutional and national guidelines for animal care.

### 2.2 Slice preparation

Hippocampal slices were prepared as described previously.^7^ Briefly, mice were anesthetized with 2–4% isoflurane inhalation and decapitated while under anesthesia. The brains were rapidly extracted and placed in ice-cold, oxygenated (95% O₂, 5% CO₂) high-Mg²⁺ dissection buffer containing the following (in mM): 130 NaCl, 24 NaHCO₃, 3.5 KCl, 1.25 NaH₂PO₄, 1 CaCl₂, 3 MgCl₂, and 10 glucose (pH 7.4). Transverse hippocampal slices (300 μm thick) were obtained using a D.S.K. Linear Slicer Pro 7 (Dosaka EM Co., Ltd., Japan). The slices were incubated in the high-Mg²⁺ dissection buffer at room temperature for at least 1 hour to allow for recovery.

Subsequently, the buffer was replaced with oxygenated artificial cerebrospinal fluid (aCSF) containing (in mM): 130 NaCl, 24 NaHCO₃, 3.5 KCl, 1.25 NaH₂PO₄, 1.5 CaCl₂, 1.5 MgCl₂, and 10 glucose (pH 7.4), and the slices were further recovered before use in intrinsic optical signal (IOS) or field recording experiments.

### 2.3 Intrinsic optical signal recording

Intrinsic optical signal recording was performed as previously described.^4,7,17^ Briefly, submerged hippocampal slices were transilluminated using a controlled infrared (IR) light source equipped with an optical filter (775 nm wavelength, Omega Filters). Images were captured using an Olympus BX50WI microscope paired with a Hamamatsu ORCA-R2 digital CCD camera. Imaging was focused on the stratum radiatum of the hippocampal CA1 region. A series of 80 images per second was acquired following a 20 Hz, 1-second electrical stimulation. The relative change in transmittance (ΔT/T) was normalized to the baseline, defined as the average transmittance of the five pre-stimulation images. The decay of the intrinsic optical signal (IOS) was calculated by averaging the last 10 seconds of the response, with responses normalized to the peak value.

### 2.4 Whole-cell recordings of long-term potentiation

Whole-cell patch-clamp recordings were conducted on pyramidal neurons within the hippocampal stratum radiatum using a Multiclamp 700B amplifier (Molecular Devices, Union City, NJ, USA). Borosilicate glass patch pipettes (resistance: 5–8 MΩ) were filled with an intracellular solution comprising (in mM): 126 potassium gluconate, 5 HEPES, 0.5 MgCl₂, and 10 BAPTA, with the pH adjusted to 7.3 using KOH. For experiments requiring astrocyte labeling, Sulforhodamine 101 (SR101) was loaded into the pipette. Positive pressure was maintained on the pipettes during advancement through the tissue. A concentric bipolar stimulation electrode (FHC, Bowdoin, ME, USA) was positioned approximately 400 μm from the patched astrocyte. Signals were low-pass filtered at 2 kHz, digitized at 10 kHz using a Digidata 1322A digitizer (Molecular Devices), and analyzed with pClamp 10.2 software (Molecular Devices). Evoked excitatory postsynaptic current (eEPSC) recordings were conducted as previously described.^7^ Briefly, eEPSCs in the CA1 stratum radiatum were induced through Schaffer collateral stimulation with a concentric bipolar electrode. Recording pipettes (resistance: 1–3 MΩ) were filled with artificial cerebrospinal fluid (aCSF). The amplitude of eEPSCs was measured and used for subsequent analyses. LTP was assessed at the CA3-CA1 synaptic pathway in the hippocampus using whole-cell patch-clamp recordings from CA1 pyramidal neurons. A stimulating electrode was positioned along the Schaffer collateral fibers to evoke eEPSCs at 0.1 Hz. LTP was induced using a theta-burst stimulation (TBS) protocol, which consisted of 10 trains of four half-maximal stimuli delivered at 100 Hz, with a 200 ms inter-train interval, while maintaining the neuron at a holding potential of 0 mV. Baseline eEPSCs were recorded for 5 minutes before TBS to ensure stability. To minimize run-up effects and optimize stimulation conditions, a test pulse (0.1 Hz) was applied during the giga-seal configuration for approximately 10 minutes, allowing fiber stabilization. Stimulation intensity was calibrated to 150–200% of the action potential threshold during this phase. LTP protocols were initiated within 10 minutes of achieving whole-cell configuration to reduce potential washout effects from the internal pipette solution. eEPSCs were recorded every 10 seconds with neurons clamped at a holding potential of −60 mV. Recording pipettes (resistance: 6–8 MΩ) were filled with an intracellular solution containing (in mM): 135 cesium methanesulfonate, 8 NaCl, 10 HEPES, 0.25 EGTA, 1 Mg-ATP, and 0.25 Na-GTP, adjusted to pH 7.2 with NaOH and an osmolarity of 290 mOsm. The eEPSC amplitudes were normalized to the average baseline amplitude for subsequent analysis.

### 2.5 Passive avoidance test

The passive avoidance test was conducted to evaluate associative learning and memory.^18^ Mice (6–7 weeks old) were placed in the light compartment of a two-chamber apparatus and allowed to explore freely for 60 seconds. After this habituation period, the door to the dark compartment was raised, and the mice were allowed to explore both compartments freely. The latency to enter the dark compartment with all four paws was recorded as the baseline latency. On day 2 (training session), the mice were again placed in the light compartment, and the latency to enter the dark compartment was recorded. Upon full entry into the dark compartment, a foot shock (0.5 mA, 2-second duration) was delivered 3 seconds after the door closed. Mice were returned to their home cages 30 seconds after the foot shock. On the test day (24 hours post-training), mice were placed back in the light compartment. After 5 seconds, the door to the dark compartment was lifted, and the latency to enter the dark compartment was recorded to assess memory retention.

### 2.6 Object-place recognition test

Behavioral testing was conducted in a gray Plexiglass box (30 × 30 × 40 cm) equipped with distinct visual cues. Animals underwent 3 days of handling before testing. To habituate to the environment, animals were placed in the arena for 10 minutes on two consecutive days. On the third day, during the training session, animals were exposed to two identical objects placed within the arena for 10 minutes. In the test session, conducted 1 hour later, animals were re-exposed to the same arena for 5 minutes. One of the objects was displaced (Displaced Object), while the other remained stationary (Stationary Object). All sessions were videotaped for subsequent analysis. Exploration behavior was assessed by experimenters blinded to the experimental groups. Exploration time was defined as the duration during which the animal oriented its head toward an object within a distance of less than 1 cm. Performance measurements were expressed as exploration ratios, calculated as follows:

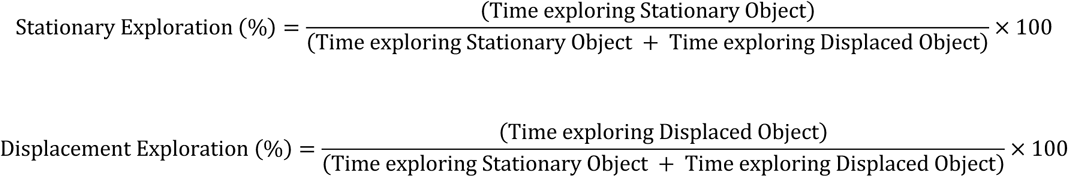

### 2.7 Stereotaxic Surgery and Viral Injection

Mice (7–8 weeks old) were anesthetized with 3–5% isoflurane inhalation and positioned in a stereotaxic frame. During surgical procedures, the isoflurane concentration was reduced to 1– 3%. Each surgery was completed within 1 hour per mouse. Viral constructs, including pSicoR lentivirus containing shRNA targeting TREK-1 (pSicoR-TREK-1-shRNA-mCherry), TRPA1 (pSicoR-TRPA1-shRNA-mCherry), and BDNF (pSicoR-BDNF-shRNA-mCherry), as well as scrambled controls (pSicoR-scrambled-shRNA-mCherry), were loaded into a microdispenser (VWR, Radnor, PA, USA) for bilateral injection into the hippocampal CA1 region (−1.7 mm AP, ± 1.7 mm ML,1.8 mm DV from the dura). A total volume of 2 μl per hemisphere was injected at a rate of 0.3 μl/min using a 25 μl syringe connected to a syringe pump (KD Scientific, USA). The stereotaxic coordinates for the injection site were 1.7 mm lateral to the bregma and 1.9 mm beneath the skull. To achieve glial-specific gene rescue, the target shRNA cassette was flanked by loxP sites to enable Cre-loxP recombination, which excised the shRNA cassette and inactivated the target shRNA.^19^ Selective retention of target gene expression in glial cells was achieved by injecting the virus into transgenic mouse lines that conditionally express Cre recombinase in glial cells, including hGFAP-CreERT2 and ALDH1/1-CreERT2.^20,21^ For experiments involving BDNF rescue, a ALDH1/1-CreERT2 mouse line crossed with BDNF floxed (BDNF fl/fl) mice was used. CreERT2 activation was induced by intraperitoneal administration of tamoxifen (1 mg dissolved in sunflower oil) or sunflower oil as a control, administered once daily for 7 consecutive days prior to shRNA injection. All experiments, including behavioral analyses and electrophysiological recordings, were performed under blinded conditions.

### 2.8 Chemicals

#### Calcium Chelation with BAPTA

BAPTA, a high-affinity calcium chelator, was utilized to disrupt intracellular calcium dynamics in astrocytes. For experiments requiring calcium chelation, BAPTA was incorporated into the intracellular solutions of glass micropipettes. The specific compositions of the internal solutions were as follows (concentrations in mM): (1) 10 potassium BAPTA and 68 potassium gluconate, (2) 40–60 potassium BAPTA, or (3) 0.1–1 potassium EGTA and 108 potassium gluconate. The osmolarity of all intracellular solutions was adjusted to 285 mOsmol using appropriate osmolarity correction methods, and the pH was titrated to 7.2 using potassium hydroxide (KOH). These preparations ensured compatibility with intracellular physiology during patch-clamp recordings.

#### Astrocyte Identification with SR101

SR101 (1 µmol/L; product code S7635, Sigma-Aldrich, St. Louis, MO, USA), a red fluorescent xanthene derivative, was employed to facilitate astrocyte identification. SR101 was included in the intracellular solution of the patch pipette at the specified concentration. Following the successful patching of an astrocyte, SR101 was allowed to diffuse through the astrocytic syncytium via gap junctions. This diffusion enabled the fluorescent labeling and subsequent visualization of the astrocyte network, thereby indicating astrocyte identity in experimental preparations.

### 2.9 Statistical analysis

Data are presented as means ± standard error mean (S.E.M.). Statistical analyses and graphing were performed using Prism 7 (GraphPad, San Jose, CA, USA) and SigmaPlot (Systat Software, San Jose, CA, USA). For comparisons between two groups, statistical significance was determined using a two-tailed Student’s t-test. For comparisons involving multiple groups, one-way or two-way analysis of variance (ANOVA) was employed. When significant interactions were detected between groups, post hoc analyses were conducted using Bonferroni, Dunnett, or multiple t-tests. In specific cases, one-way ANOVA followed by Bonferroni, Dunnett, or Tukey post hoc tests was applied to identify homogeneous subsets.

## 3. Results

### 3.1 Astrocytic TREK-1 is essential for synaptic plasticity and spatial memory

To investigate the relationship between astrocytic volume transients and synaptic plasticity, we employed the intrinsic optical signal (IOS) imaging technique^17,22,23^ in order to indirectly observe transient volume changes in real-time from hippocampal slices by detecting light transmittance during intense neuronal activity. The neuronal activity-induced volume transient was defined as the astrocytic volume change occurring within one minute of intense neuronal activity. IOS imaging and electrophysiological recordings were conducted simultaneously in CA1 neurons of the hippocampus (Fig. 1A) Stimulating electrodes were positioned in the stratum radiatum of the CA3-CA1 Schaffer collateral pathway, enabling precise correlation between light transmittance changes and synaptic plasticity. LTP was induced using a theta-burst stimulation (TBS) protocol, which consisted of 10 trains of four half-maximal stimuli delivered at 100 Hz, with a 200 ms inter-train interval, while maintaining the neuron at a holding potential of 0 mV. Baseline eEPSCs were recorded for 5 minutes before TBS to ensure stability. TBS elicited a rapid increase in change of light transmittance, accompanied by a simultaneous increase in normalized amplitude of eEPSC, occurring within seconds (Fig. 1B). During short-term potentiation (STP), the normalized amplitude of eEPSC exhibited a decline (exponential decay kinetics) over 10 minutes before stabilizing into long-term potentiation at over 150% of baseline, while light transmittance returned to baseline within the same timeframe. These results indicate that the IOS changes mirror the dynamics of STP, with both signals showing a temporally correlating reduction following TBS.

**Figure 1.**
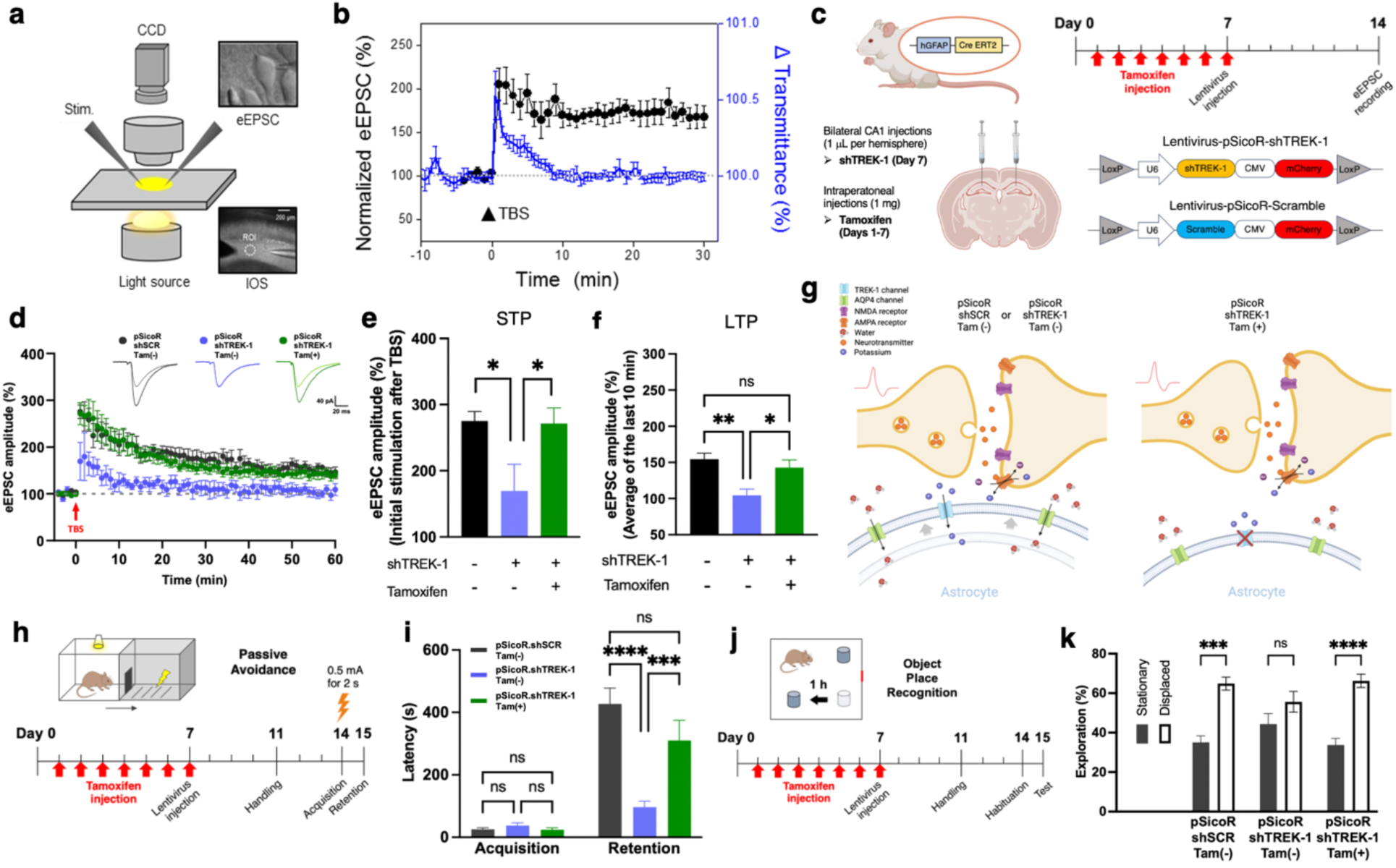
Astrocytic TREK-1 is essential for synaptic plasticity and spatial memory. **(a)** Schematic representation of the IOS imaging and electrophysiological setup. The CA1 region of hippocampal slices was illuminated using an infrared light source, and changes in transmittance (ΔT/T) were recorded. eEPSCs were simultaneously measured using whole-cell patch-clamp techniques. **(b)** IOS and eEPSC responses to TBS. TBS induces a rapid transient decrease in IOS (blue) corresponding with a significant increase in normalized eEPSC amplitudes (black). These changes reflect activity-dependent synaptic modifications. Data represent mean ± SEM. **(c)** Experimental design for astrocyte-specific TREK-1 knockdown and rescue. GFAP-Cre.ERT2 transgenic mice received bilateral CA1 injections of lentivirus encoding shTREK-1 or scrambled shRNA (control). Tamoxifen was administered for 7 days to induce astrocyte-specific TREK-1 knockdown, followed by electrophysiological assays 7 days post-injection. **(d)** Representative traces and quantification of eEPSC amplitude during 60 minutes post-TBS. TREK-1-deficient mice (pSicoR-shTREK-1 Tam(-)) exhibit significantly impaired short-term potentiation (STP) and long-term potentiation (LTP) compared to controls (pSicoR-shSCR Tam(-)). TREK-1 rescue (pSicoR-shTREK-1 Tam(+)) restores normal synaptic plasticity. **(e, f)** Quantification of eEPSC amplitudes immediately after TBS **(e)** and at the final 10 minutes of recording **(f)**. TREK-1 knockdown reduces synaptic potentiation, which is rescued by TREK-1 re-expression. Statistical comparisons: one-way ANOVA with post hoc tests (*p < 0.05; **p < 0.01; ns = not significant). **(g)** Mechanistic model for TREK-1-mediated regulation of astrocytic volume transient. TREK-1 channels mediate potassium influx during synaptic activity, creating osmotic gradients that facilitate water influx via AQP4. The resulting astrocytic swelling may mechanically modulate synapse function. In TREK-1-deficient astrocytes, impaired potassium buffering disrupts these processes, reducing synaptic efficacy. **(h-k)** Behavioral consequences of TREK-1 deficiency. **(h)** Passive avoidance task timeline. **(i)** TREK-1-deficient mice exhibit reduced retention latency, indicative of long-term memory deficits. **(j)** Object-place recognition test timeline. **(k)** TREK-1-deficient mice display reduced exploration of displaced objects, indicating spatial memory impairments. TREK-1 rescue restores normal performance. Statistical comparisons: two-way ANOVA with post hoc tests (***p < 0.001; ****p < 0.0001; ns = not significant). Data represent mean ± SEM.

To explore the role of astrocytic TREK-1 in synaptic plasticity, we employed GFAP-CreERT2 transgenic mice for astrocyte-specific gene silencing using a tamoxifen-inducible Cre-lox system (Fig. 1C). Mice received tamoxifen injections for 7 days, followed by bilateral CA1 injections of lentivirus encoding either shTREK-1_mCherry or control shSCR_mCherry. Seven days after viral delivery, hippocampal slices were prepared for electrophysiological recordings. In control mice (pSicoR-shSCR Tam (-)), TBS reliably induced a robust STP and LTP, evidenced by sustained increases in normalized eEPSC amplitudes (Fig. 1D-1F). In contrast, TREK-1 gene-silencing (pSicoR-shTREK-1 Tam (-)) significantly impaired both STP and LTP, as shown by a diminished eEPSC peak and a gradual return toward baseline. Importantly, astrocytic TREK-1 rescue (pSicoR-shTREK-1 Tam (+)) restored STP and LTP, with normalized eEPSC amplitudes comparable to controls (Fig. 1E and 1F). Quantitative analysis showed a significant reduction in post-TBS eEPSC amplitudes in TREK-1-deficient mice compared to both control and astrocytic TREK-1-rescue groups, indicate the necessity of astrocytic TREK-1 for synaptic plasticity.

A mechanistic model for astrocytic TREK-1 function in synaptic plasticity involves potassium buffering during synaptic activity (Fig. 1G). TREK-1 channels mediate astrocytic potassium efflux, preventing extracellular potassium accumulation that could impair neuronal function. This efflux generates an osmotic gradient, facilitating water influx through astrocytic aquaporin-4 (AQP4) channels. The resulting astrocytic swelling may influence synaptic plasticity through mechanical signaling at neuron-astrocyte interfaces. In TREK-1-deficient astrocytes, impaired potassium buffering likely disrupts these processes, leading to reduced synaptic plasticity.^24^

We further examined the behavioral consequences of TREK-1 deficiency using passive avoidance and object place recognition tests (Fig. 1H and 1J). TREK-1-deficient mice exhibited impaired retention in the passive avoidance task compared to controls, as reflected by significantly reduced latency during the retention phase (Fig. 1I). Similarly, in the object place recognition task, TREK-1-deficient mice showed reduced exploration of displaced objects, indicating deficits in spatial memory (Fig. 1K). Astrocytic TREK-1 rescue restored performance in both tasks to levels comparable to controls, suggesting a critical function of astrocytic TREK-1 in spatial memory formation. Together, these findings depict an essential role of astrocytic TREK-1 in synaptic plasticity and memory, facilitated through its TREK-1-mediated volume transient function during potassium buffering.

### 3.2 Astrocytic TRPA1 deficiency disrupts synaptic plasticity and memory formation

To explore the potential role of astrocytic volume transient-induced calcium signaling in synaptic plasticity and memory, we investigated TRPA1 channels due to their mechanosensitive characteristics and function in astrocytic calcium signaling. The role of astrocytic TRPA1 in synaptic plasticity was examined through electrophysiological recordings of eEPSCs in hippocampal slices obtained from TRPA1 WT and KO mice. Following TBS, TRPA1 WT mice displayed strong LTP, evident in a sustained elevated eEPSC amplitude over 60 minutes (Fig. 2A). In contrast, TRPA1 KO mice showed markedly impaired LTP compared to TRPA1 WT mice, with eEPSC amplitudes declining to baseline by the end of the recording period (Fig. 2A). STP was assessed and found to remain unchanged between TRPA1 KO and WT mice (Fig. 2B), as indicated by comparable eEPSC amplitudes immediately following TBS. This finding suggests that TRPA1 deficiency specifically affects long-term synaptic plasticity rather than STP mechanisms. Quantitative analysis further demonstrated the significant differences in LTP between TRPA1 WT and KO mice. eEPSC amplitudes were significantly lower in TRPA1 KO mice compared to TRPA1 WT mice during the sustained phase of LTP (Fig. 2C). These results collectively indicate that astrocytic membrane stretching during the astrocytic volume transient process activates the mechanosensitive function of TRPA1 channels, contributing to their involvement in LTP. Furthermore, the reduction in eEPSC amplitudes observed in LTP but not in STP in TRPA1 KO mice suggests that STP is upstream of the involvement of TRPA1 in synaptic plasticity, emphasizing a specific role for TRPA1 in the later stages of synaptic potentiation.

**Figure 2.**
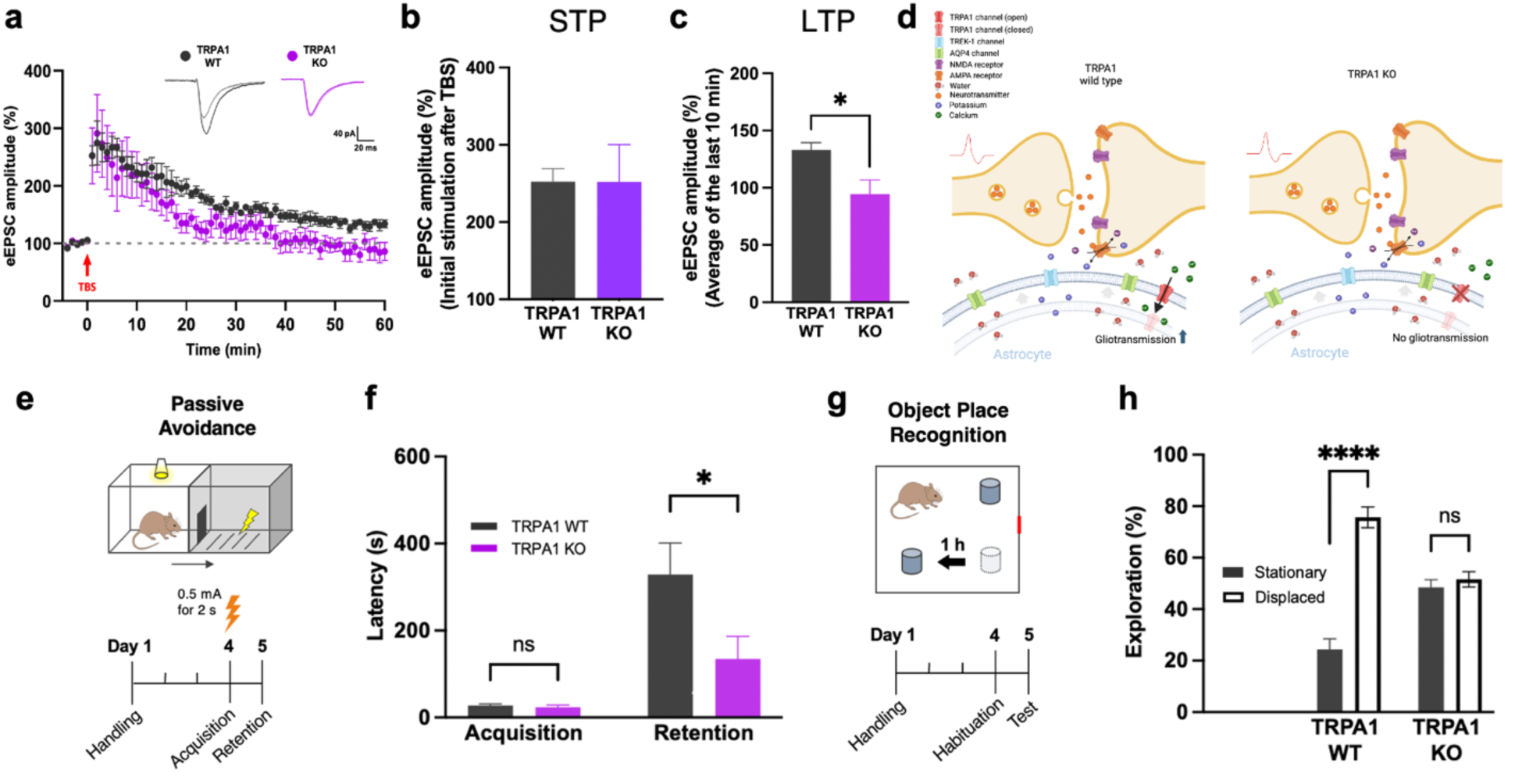
Astrocytic TRPA1 is essential for synaptic plasticity and spatial memory. **(a)** Time-course of eEPSC amplitudes recorded from hippocampal CA1 neurons following TBS. TRPA1 WT mice exhibit sustained LTP over 60 minutes, whereas TRPA1 KO mice show impaired LTP with eEPSC amplitudes returning to baseline. Representative traces are shown above the graph for TRPA1 WT (black) and TRPA1 KO (purple). Data are presented as mean ± SEM. **(b, c)** Quantification of eEPSC amplitudes immediately after TBS **(b)** and during the final 10 minutes of recording **(c)**. TRPA1 KO mice exhibit significantly reduced eEPSC amplitudes compared to TRPA1 WT mice, indicating a critical role for TRPA1 in maintaining LTP. Statistical analysis: Student’s t-test (*p < 0.05; ns = not significant). **(d)** Schematic model illustrating the role of TRPA1 in astrocyte-neuron signaling. In TRPA1 WT astrocytes, potassium and water influx during synaptic activity activate mechanosensitive TRPA1 channels, leading to calcium influx and gliotransmitter release. This enhances synaptic efficacy. In TRPA1 KO astrocytes, the absence of calcium signaling prevents gliotransmitter release, impairing synaptic plasticity. **(e-h)** Behavioral analyses reveal deficits in long-term memory in TRPA1 KO mice. **(e)** Passive avoidance test timeline. **(f)** TRPA1 KO mice exhibit significantly reduced retention latency compared to TRPA1 WT mice, indicating impaired associative memory. **(g)** Object-place recognition test timeline. **(h)** TRPA1 KO mice display reduced exploration of displaced objects, reflecting deficits in spatial memory. Statistical analysis: two-way ANOVA with post hoc tests (*p < 0.05; ****p < 0.0001; ns = not significant). Data are presented as mean ± SEM.

Mechanistically, astrocytic TRPA1 channels appear to contribute to synaptic plasticity by responding to astrocytic swelling induced by neuronal activity. Potassium and water influx in astrocytes during synaptic activity activate TRPA1 channels via mechanosensitive gating, initiating calcium influx and subsequent gliotransmitter release (Fig. 2D). The absence of TRPA1-mediated calcium signaling in KO mice likely interrupts critical astrocyte-neuron communication, impairing LTP.

The behavioral relevance of TRPA1 deficiency was evaluated using the passive avoidance and object place recognition tasks. TRPA1 KO mice exhibited significantly shorter retention latencies in the passive avoidance test compared to WT mice, indicating long-term memory deficits (Fig. 2E, 2F). Similarly, TRPA1 KO mice showed reduced exploration of displaced objects in the object place recognition test, indicative of spatial memory impairments (Fig. 2G, 2H). These results uncover an essential role of astrocytic TRPA1 in maintaining synaptic plasticity and supporting memory functions through its involvement in calcium signaling.

### 3.3 Astrocytic calcium and BDNF as determinants of synaptic plasticity and memory

To elucidate the role of astrocytic signaling in synaptic plasticity, we adopted the necessity and sufficiency experimental framework, modeled after the approach used in a previous study^25^, where astrocytic calcium was clamped, and plasticity was subsequently rescued with a gliotransmitter. Following this paradigm, we sought to demonstrate that astrocytic calcium signaling is necessary for synaptic plasticity. To establish necessity, astrocytes within the CA1 of hippocampal slices from B6-GFAP-GFP mice were patched with micropipettes loaded with the high-affinity calcium chelator BAPTA and SR101, a red fluorescent xanthene dye that diffuses through gap junctions (Fig. 3A). SR101 diffusion enabled clear visualization of the astrocyte syncytium, ensuring astrocyte specificity of the calcium clamp. BAPTA sequestered intracellular calcium, effectively clamping astrocytic calcium dynamics, as previously described.^25^ In BAPTA-loaded syncytial astrocyte slices, TBS failed to induce STP or LTP, as indicated by eEPSC amplitudes not significantly exceeding baseline value following TBS (Fig. 3B-3D).

**Figure 3.**
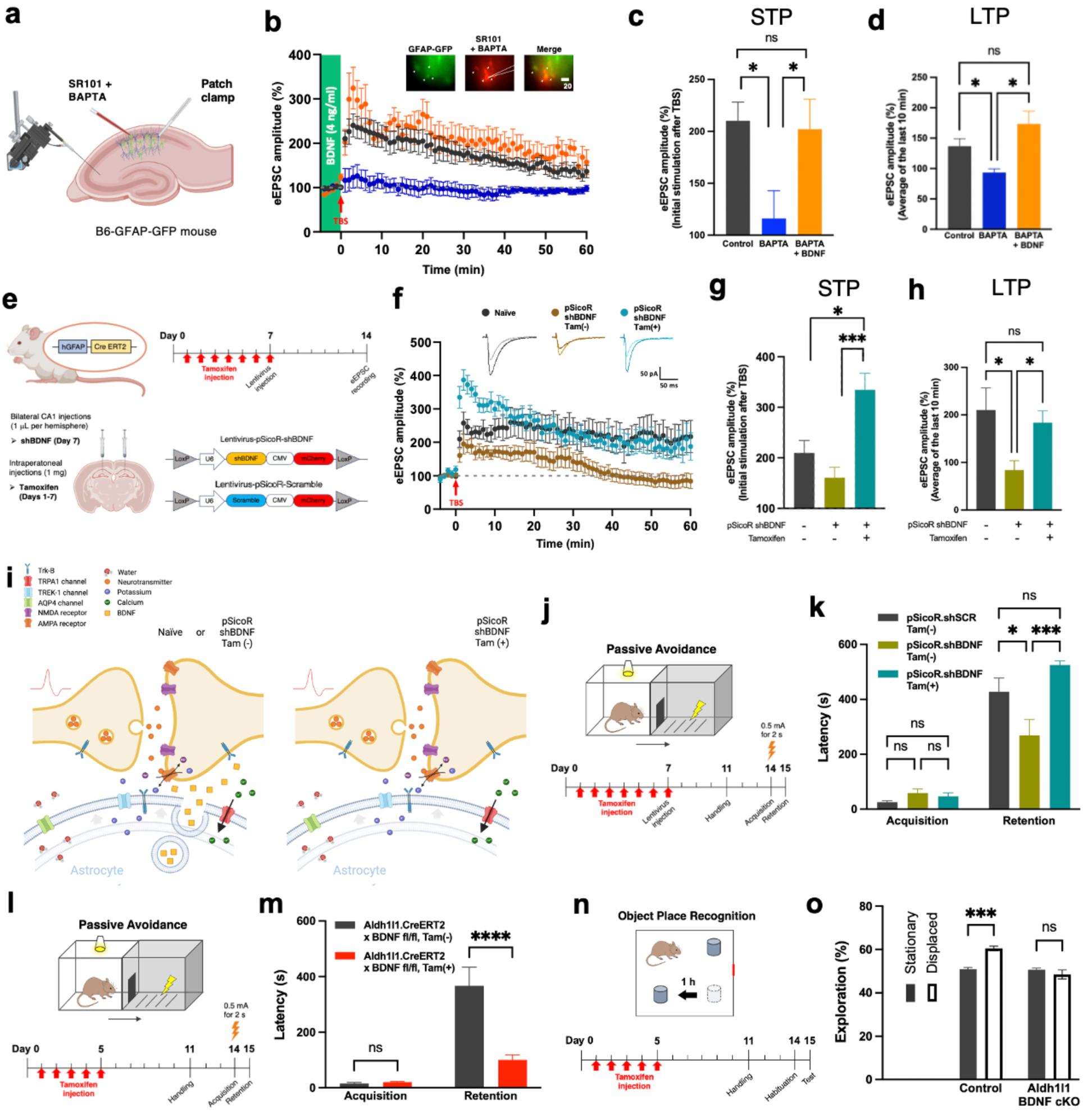
Astrocytic BDNF supports synaptic plasticity and memory by modulating calcium dynamics. **(a)** Schematic of hippocampal slice preparation and treatment with BAPTA and SR101. Astrocytes were loaded with BAPTA to chelate intracellular calcium, and SR101 dye was used for astrocyte visualization. **(b)** Time-course of eEPSC amplitudes following TBS in hippocampal slices. Control slices show robust LTP, while BAPTA-treated slices exhibit impaired LTP. Supplementing BAPTA-treated slices with BDNF restored LTP. Insets show astrocytic SR101 labeling and BAPTA loading. Data are presented as mean ± SEM. **(c, d)** Quantification of eEPSC amplitudes immediately after TBS **(c)** and during the final 10 minutes of recording **(d)**. BAPTA treatment reduces synaptic potentiation, which is rescued by BDNF supplementation. Statistical comparisons: one-way ANOVA with post hoc tests (*p < 0.05; ns = not significant). **(e)** Experimental timeline for astrocyte-specific knockdown and rescue of BDNF. Mice received bilateral CA1 injections of lentivirus encoding shBDNF or scrambled control. Astrocytic BDNF expression was restored via tamoxifen treatment. **(f)** eEPSC amplitude time-course following TBS in naïve, shBDNF, and tamoxifen-treated shBDNF mice. BDNF knockdown impairs LTP, while tamoxifen-mediated rescue restores normal synaptic plasticity. Representative traces are shown above the graph. **(g, h)** Quantification of eEPSC amplitudes immediately after TBS **(g)** and during the final 10 minutes of recording **(h)**. Statistical analysis confirms significant rescue of LTP in tamoxifen-treated shBDNF mice (*p < 0.05; **p < 0.01; ns = not significant). **(i)** Schematic model illustrating the role of astrocytic BDNF in synaptic plasticity. Calcium influx through TRPA1 channels in astrocytes triggers BDNF release, enhancing presynaptic neurotransmitter release. In BDNF-deficient astrocytes, this pathway is disrupted, impairing synaptic efficacy and plasticity. **(j-o)** Behavioral assessments of BDNF function in memory. **(j, l)** Passive avoidance task timelines. **(k, m)** BDNF knockdown reduces retention latency, reflecting deficits in long-term memory, which are rescued by tamoxifen-mediated restoration of astrocytic BDNF. **(n)** Object-place recognition test timeline. **(o)** Exploration ratios during object-place recognition. BDNF knockdown impairs spatial memory, as indicated by reduced exploration of displaced objects. Tamoxifen treatment restores performance. Statistical analysis: two-way ANOVA with post hoc tests (***p < 0.001; ****p < 0.0001; ns = not significant). Data are presented as mean ± SEM.

These findings demonstrate that astrocytic volume transient-induced calcium signaling is indispensable for mediating synaptic plasticity. To assess whether astrocytic calcium signaling modulates synaptic plasticity via the release of BDNF, we examined whether BDNF alone is sufficient to restore synaptic plasticity in conditions of calcium depletion, given its well-established roles in synaptic plasticity and memory.^8^ Sufficiency was assessed by introducing exogenous BDNF into the same experimental setup (Fig. 3B). Exogenous BDNF treatment in BAPTA-treated slices fully restored STP and LTP, elevating eEPSC amplitudes during the final 10 minutes of recording to levels comparable to untreated controls (Fig. 3C and 3D). These results indicate that BDNF is sufficient to compensate for the loss of calcium-dependent astrocytic signaling.

To further explore the necessity of astrocytic BDNF, GFAP-CreERT2 mice were utilized to conditionally silence general BDNF expression. Mice received tamoxifen injections to activate Cre, followed by bilateral CA1 lentiviral injections of shBDNF (Fig. 3E). In slices from these animals, TBS-induced LTP was significantly impaired, with eEPSC amplitudes rapidly declining post-TBS (Fig. 3F). STP, however, was unaffected (Fig. 3G), indicating that astrocytic BDNF specifically contributes to sustained synaptic plasticity. Restoration of astrocytic BDNF expression via tamoxifen rescue significantly increased LTP (Fig. 3H), proposing astrocytic BDNF as a critical downstream effector of calcium signaling.

These findings demonstrate that astrocytic calcium signaling is essential for gliotransmitter release, with astrocytic BDNF serving as both a necessary and sufficient mediator of LTP. This expands on the necessity and sufficiency framework established for D-serine in synaptic plasticity, implicating BDNF as a pivotal component of astrocytic regulation of LTP. Mechanistically, BDNF release is driven by calcium influx through mechanosensitive TRPA1 channel activation, triggered by astrocytic swelling during potassium and water uptake, which may enhance synaptic plasticity through pre-, post-, and peri-synaptic activation of tyrosine receptor kinase B (Trk-B) (Fig. 3I).

To assess the impact of astrocytic BDNF deficiency on memory, we performed passive avoidance (Fig. 3J–3M) and object place recognition tests (Fig. 3N and 3O). In the passive avoidance task, we used GFAP-CreERT2 transgenic mice to induce general BDNF gene-silencing or rescue with tamoxifen (Fig. 3J). These behavioral assays revealed significant deficits in both spatial and long-term memory. Mice lacking BDNF exhibited reduced retention latency, indicative of impaired long-term memory consolidation (Fig. 3K). Astrocytic BDNF rescue via tamoxifen administration restored retention latency to durations similar to controls. To further explore whether these effects extended beyond the GFAP-positive astrocyte population, as well as to determine the effects of astrocyte-specific gene-silencing of BDNF, we utilized ALDH1/1-CreERT2 x BDNF fl/fl mice, which target a broader subset of astrocytes (Fig. 3L).^15,26^ Tamoxifen-mediated removal of astrocytic BDNF led to significant retention deficits, demonstrating the necessity of astrocytic BDNF in spatial memory (Fig. 3M). The object place recognition test was utilized to further validated the necessity of BDNF for long-term memory (Fig. 3N). In control mice, a greater proportion of time was spent exploring displaced objects, reflecting intact spatial memory (Fig. 3O). In contrast, ALDH1/1-CreERT2 x BDNF fl/fl mice with tamoxifen-induced astrocytic BDNF deficiency showed impaired spatial memory, as indicated by diminished exploration of displaced objects. Together, these findings demonstrate an integral role of astrocytic BDNF in synaptic plasticity and memory processes, possibly mediated through astrocytic volume transient- and calcium-dependent mechanisms.

## 4. Discussion

This study elucidates the critical role of astrocytic volume transients in regulating synaptic plasticity and memory formation, with TREK-1-mediated potassium buffering, TRPA1-dependent calcium signaling, and astrocytic BDNF release emerging as key components of this process. By integrating these findings with prior research^4,7,25^, we provide a cohesive framework highlighting astrocytic contributions to synaptic plasticity and cognitive functions. We propose a comprehensive model in which 1) astrocytic TREK-1 channels initiate potassium buffering and volume transients, 2) TRPA1 channels convert these mechanical signals into calcium influx, and 3) the resulting astrocytic release of BDNF enhances synaptic plasticity. The dynamic interplay between synaptic activity and astrocytic volume transients alludes to their role in synaptic plasticity. Our observations of synchronous increases in IOS changes and normalized eEPSC amplitudes following TBS (Fig. 1B) illustrate the precise coupling between astrocytic responses and synaptic plasticity. The rapid rise and subsequent normalization of IOS signals, mirroring the decay kinetics of STP, suggest that astrocytic volume transients actively contribute to synaptic potentiation. This mechanism facilitates sustained elevation in eEPSC amplitudes that stabilize at over 50% of the baseline levels during LTP.

Our findings demonstrate that TREK-1 plays a pivotal role in modulating both STP and LTP, as evidenced by significant impairments in these processes following TREK-1 gene-silencing, accompanied by deficits in spatial and long-term memory tasks, underscoring the importance of TREK-1 in astrocytic volume dynamics and, consequently, synaptic plasticity and memory. In contrast, our data indicate that TRPA1 channels, while essential for LTP, do not influence STP, suggesting a TREK-1-specific mechanism in the initial phases of synaptic potentiation. Recent findings by Woo et al. provide compelling support for this conclusion by identifying TREK-1 as a mediator of fast, calcium-independent glutamate release from astrocytes.^21^ This mode of release is initiated through the activation of G_αi_-protein coupled receptor signaling and the dissociation of G_βγ_ subunits, which directly interact with TREK-1 to open its channel to release glutamate.^21^ It is possible that this rapid glutamate release could target metabotropic glutamate receptors facilitating transient synaptic enhancement characteristic of STP. However, this possibility has not been tested in this study. Because TREK-1-mediated release of glutamate requires activation of G_αi_-protein coupled receptor signaling, the TREK-1 mediated volume transients may not involve glutamate release through TREK-1. These possibilities require future investigations.

Our findings demonstrate that TRPA1 deficiency impairs LTP without affecting STP, indicating that TRPA1-mediated calcium signaling is essential for LTP, but not STP. Similarly, BDNF gene-silencing disrupts LTP while sparing STP, further supporting that STP is regulated by mechanisms independent of TRPA1-mediated astrocytic calcium signaling. These findings, distinct mechanisms for STP and LTP, emphasize the role of astrocytic volume dynamics in regulating STP. Recent work conducted by Ucar et al., 2021 and Kasai et al., 2023 attribute STP to mechanical transmission initiated by post-synaptic spine swelling following neuronal activity, which causes bumping into the presynaptic neuron, thereby increasing presynaptic neurotransmitter release through enhanced SNARE assembly.^27,28^ Based on our study, we propose an alternative in which astrocytic volume transients may underlie mechanical transmission in STP through astrocytic bumping of the pre-synaptic membrane. This proposed idea is supported by our findings that TREK-1 gene-silencing—known to impair astrocytic volume changes^4^—reduces STP, while TREK-1 restoration rescues it. This exciting possibility warrants further investigation.

The foundational work by Henneberger et al. demonstrated that astrocytic calcium signaling is necessary for synaptic plasticity, largely through the release of D-serine as a gliotransmitter to activate NMDA receptor co-agonist sites.^25^ While this work underscored D-serine’s importance in LTP, emerging evidence^29,30^—including findings from our investigation— suggests that astrocytic BDNF serves as a more versatile and potent modulator of LTP. Recent research, such as that of Koh et al., demonstrated that astrocytic D-serine contributes predominantly to heterosynaptic long-term depression (LTD), a process integral to cognitive flexibility.^30^ These findings, while highlighting D-serine’s role in synaptic plasticity, reveal its relatively specialized function in heterosynaptic LTD but not in homosynaptic LTP, contrasting with the broader and multifaceted effects of astrocytic BDNF. Unlike D-serine, which acts primarily by enhancing post-synaptic NMDAR activity, astrocytic BDNF exerts influence across pre-, post-, and peri-synaptic membranes via TrkB receptors.^8,14,31^ Our study demonstrates that BDNF is both necessary for LTP induction and sufficient for LTP restoration under conditions of disrupted astrocytic calcium dynamics, as evidenced by experiments involving BAPTA-mediated calcium clamping and targeted astrocytic gene-silencing. Importantly, behavioral assays revealed significant impairments in memory and plasticity following astrocyte-specific BDNF gene-silencing, further supporting its critical role in spatial and long-term memory. The work by Koh et al. reinforces our findings by elucidating the mechanism by which astrocytic D-serine release regulates NMDAR tone and LTD.^30^ However, their data also underscore the limitations of D-serine as a modulator of plasticity, given its confined role in regulating NMDAR-dependent LTD rather than LTP. Collectively, these results suggest that astrocytic BDNF, with its capacity to modulate diverse synaptic targets and restore LTP under impaired calcium signaling, represents a more comprehensive gliotransmitter for LTP than D-serine.

While our findings highlight the roles of TREK-1-mediated astrocytic volume transients and TRPA1-dependent BDNF release in synaptic plasticity, future studies leveraging advanced optogenetic tools, such as Opto-Stim1^32^, could enable more precise astrocyte-specific calcium modulation, offering a system to directly investigate the interplay between astrocytic calcium signaling, BDNF release, and memory. In addition, while we successfully identified behavioral deficits linked to impaired LTP in long-term memory tasks, we did not explore the functional implications of impaired STP on short-term memory. Implementation of short-term memory tasks, such as Y-maze alternation and brief-interval novel object recognition, could shed light on the specific contributions of astrocytic volume transients to short-term memory.

In conclusion, this study establishes astrocytic volume transients as integral components of synaptic plasticity and memory. By elucidating the roles of TREK-1, TRPA1, and BDNF, we provide a comprehensive framework for understanding astrocyte-neuron interactions via physical volume changes and their impact on cognition. These findings lay the groundwork for novel therapeutic approaches targeting astrocytic volume pathways to enhance synaptic plasticity and memory.

## Author Contributions

C.J.L. conceived and designed this project. J.W., J.M.L., and JH.W. performed the experiments. J.W., J.M.L., and V.J.D. analyzed the data. V.J.D. wrote the manuscript with inputs from other authors. C.J.L. edited the manuscript.

## Acknowledgments

We would like to that the Center for Cognition and Sociality (IBS-R001-D2) under the Institute for Basic Science (IBS), Republic of Korea for funding this study. We acknowledge the use of ChatGPT 4.0 for assistance with writing style, outlining, and proofreading. All content presented in this manuscript is original. The authors also extend acknowledgement to BioRender.com for enabling the creation of images.

## Conflicts of Interest

The authors declare no conflicts of interest.

## Data Availability Statement

The data that support the findings of this study are available on request from the corresponding author.

## Notes

### Competing Interest Statement

The authors have declared no competing interest.

